# Apollo-IRE1: A Genetically Encoded Sensor for Live Cell and Multiplexed Imaging of ER Stress

**DOI:** 10.64898/2026.03.20.712661

**Authors:** Eric J. Floro, Alex M. Bennett, Romario Regeenes, Huntley H. Chang, Nitya Gulati, Kenneth K.Y. Ting, Jonathan V. Rocheleau

**Affiliations:** Institute of Biomedical Engineering, University of Toronto, Toronto, M5S 3G9, Canada; Toronto General Hospital Research Institute, University Health Network, Toronto, M5G 2C4, Canada; Department of Physiology, University of Toronto, Toronto, M5S 1AB, Canada; Banting and Best Diabetes Centre, University of Toronto, Toronto, M5G 2C4, Canada

**Keywords:** fluorescence anisotropy, homoFRET, live-cell imaging, unfolded protein response (UPR), IRE1α, ER stress, pancreatic beta cells, protein oligomerization, BiP/GRP78, TXNIP

## Abstract

Pancreatic beta cells face exceptional protein folding demands from high insulin production requirements, placing extraordinary stress on the ER and contributing to dysfunction in diabetes pathogenesis. Monitoring ER stress dynamics in living cells remains challenging due to the destructive nature of traditional biochemical methods and the limitations of existing fluorescent sensors. Here, we present Apollo-IRE1, a genetically encoded sensor that reports on stress-induced IRE1 oligomerization and associated change in homoFRET via changes in fluorescence anisotropy. Apollo-IRE1 provides a ratiometric, intensity-independent readout, resulting in low day-to-day variability and a minimal spectral bandwidth, enabling multiplexed imaging alongside other cellular parameters. Photobleaching and enhancement curve analysis show that Apollo-IRE1 exists in apparent monomeric, dimeric, and oligomeric states corresponding to baseline, moderate, and terminal ER stress conditions. The sensor also responds rapidly to chemical and physiological ER stressors in both immortalized beta-cell lines and primary mouse islet cells. These data establish Apollo-IRE1 as a practical tool for investigating ER stress dynamics in beta cells and other contexts where longitudinal single-cell measurements are essential.

---

Pancreatic beta cells are specialized insulin-producing endocrine cells located within the islets of Langerhans. Beta-cell insulin production and secretion are essential for maintaining normal blood glucose levels, as insulin stimulates glucose uptake by the liver, muscle, and adipose tissue. An individual beta cell can produce up to 1 million molecules of insulin per minute, with proinsulin accounting for up to 50% of total protein synthesis under stimulated conditions.^1,2^Given that insulin contains three intrachain disulphide bonds, beta cells must therefore coordinate the formation of approximately 3 million disulphide bonds per minute.^1^ This immense protein folding load places exceptional demands on the endoplasmic reticulum (ER). Chronic overload of ER capacity is now widely appreciated to contribute to beta-cell dysfunction associated with type 2 diabetes (T2D), and thus a potential druggable target for treating the disease.^3–5^

Protein folding capacity in the ER is monitored and mediated by three transmembrane sensors: inositol-requiring enzyme 1 (IRE1), PKR-like ER-resident kinase (PERK), and activating transcription factor 6 (ATF6). IRE1 is the most evolutionarily conserved receptor that senses protein misfolding in the ER lumen and signals through dual-function kinase/RNase activity to trigger the unfolded protein response (UPR).^6–8^ The UPR works to restore ER homeostasis by increasing the transcription of ER chaperone proteins, enabling ER-associated degradation (ERAD) of misfolded proteins, and transiently reducing protein translation to relieve ER protein load.^9–11^ If these protective steps fail to restore homeostasis, the UPR transitions from adaptive to terminal, triggering beta-cell apoptosis.^12,13^

BiP (GRP78) is an ER chaperone that binds IRE1 under basal conditions and dissociates during ER stress as misfolded proteins accumulate.^14^ Upon BiP dissociation, IRE1 autophosphorylates its kinase domain, thereby activating RNase activity.^15^ This activity splices XBP1 (X-box-binding protein 1) into a transcription factor that triggers adaptive functions, making spliced XBP1 a key marker of early adaptive UPR.^16,17^ Chronic, unresolved UPR results in increased regulated IRE1-dependent decay (RIDD) and higher-order oligomerization of IRE1.^18–20^ The transition to this terminal state is also associated with the induction and activation of thioredoxin-interacting protein (TXNIP), a multifunctional protein that activates the NLRP3 inflammasome and induces cell death.^9,21^ Thus, we aimed to develop a sensor capable of tracking the transition from BiP-mediated adaptive UPR to terminal UPR.

Current methods for measuring ER stress and UPR activation are endpoint assays that rely on sample destruction/fixation prior to measurement (e.g., immunoblotting and western blotting to detect the phosphorylation levels of IRE1α and PERK, or the cleavage of ATF6α),^22^ which impede tracking the dynamic progression of ER stress. Additionally, these methods lack sufficient spatial and temporal resolution to follow the responses of specific cells within a tissue (e.g., beta cells within islets) and to capture transient changes in ER stress signaling.^22^ Recent work has explored translating ER stress sensors into fluorescent reporters for live-cell experiments. To monitor the IRE1 branch of the UPR, the Walter lab developed constructs of human IRE1α tagged with intersequence fluorescent proteins (e.g., EGFP and mCherry) to measure IRE1 oligomerization by heteroFRET.^23,24^ While this strategy was effective at confirming the molecular oligomerization of IRE1, heteroFRET sensors that require the expression of two fluorescent proteins are inherently unreliable due to normal variation in the ratio of the two constructs (day-to-day and cell-to-cell) and consume a significant portion of the visible spectrum, limiting their use for multiplexing the study of ER stress and metabolism in the same cell.^23,25–27^

We and others previously reported genetically encoded sensors based on homoFRET (i.e., FRET between identical fluorescent proteins), with changes in sensor oligomerization detected by steady-state fluorescence anisotropy (i.e., emission polarization).^26,28^ Steady-state fluorescence anisotropy is independent of illumination power and sensor concentration, and varies little between instruments, making it valuable for high-content imaging. In this approach, the single fluorescent protein is also swappable, thus facilitating multispectral imaging with other sensors.^26,28,29^ Our first sensor (Apollo-NADP^+^) was based on NADP^+^-dependent oligomerization of a mutated form of glucose-6-phosphate dehydrogenase.^26,30–32^ To construct an Apollo-IRE1 sensor, we here explore an IF2L mutant of IRE1 (WLLI359-362 to GSGS359-362) tagged with intrasequence GFP that was previously generated by the Walter lab.^24^ The IF2L mutation disrupts the lumenal domain oligomerization interface while preserving the IF1L dimerization interface, restricting the sensor to report on dimerization rather than higher-order oligomerization. Notably, the IF2L mutant lacks XBP1 splicing activity when expressed in IRE1-knockout cells.^24^ However, in cells with endogenous IRE1, XBP1 splicing proceeds normally, allowing the sensor to be used without disrupting native UPR signaling. The construct was also previously used to measure IRE1 oligomerization by heteroFRET.^24^

## RESULTS

### Apollo-IRE1 Sensor Design

The Walter lab previously created constructs of human IRE1 (‘IF2L mutant’) tagged with intrasequence fluorescent proteins (e.g., EGFP and mCherry) to measure IRE1α oligomerization by heteroFRET.^24^HeteroFRET consumes a significant portion of the visible spectrum, making it challenging to implement multiplex imaging. To assess whether these constructs could be adapted for a single-color Apollo sensor, we replaced EGFP with mVenus in the IF2L mutant construct to create the “Apollo-IRE1” sensor (**Figure 1a**). We postulated that a progressive decrease in fluorescence anisotropy of the mVenus construct would reflect stress-induced changes in IRE1 oligomerization, from an inactive baseline to activated states. To validate Apollo-IRE1, we first confirmed that INS1E beta cells expressing the sensor continued to splice XBP1 as expected (**Figure S1a**). We then expressed the sensor in INS1E cells and imaged the fluorescence anisotropy response after treatment with common inducers of ER stress, including thapsigargin (Tg) to deplete the ER of Ca^2+^ and dithiothreitol (DTT) to reduce disulphide bonds in the ER (**Figure 1b–c**). Apollo-IRE1 was consistently absent from the nucleus and showed a web-like feature consistent with retention in the ER (**Figure 1b, top**). The representative anisotropy images show a drop in anisotropy across the cells after treatment with Tg (12 h) and DTT (3 h). Measuring these responses across many cells (*N*) and multiple experimental days (*n*) showed consistent drops in fluorescence anisotropy from control cells, with the most significant drop found after DTT treatment (**Figure 1c**). These data suggest that the single-color Apollo-IRE1 sensor can be used to monitor ER stress in living cells.

**Figure 1.**
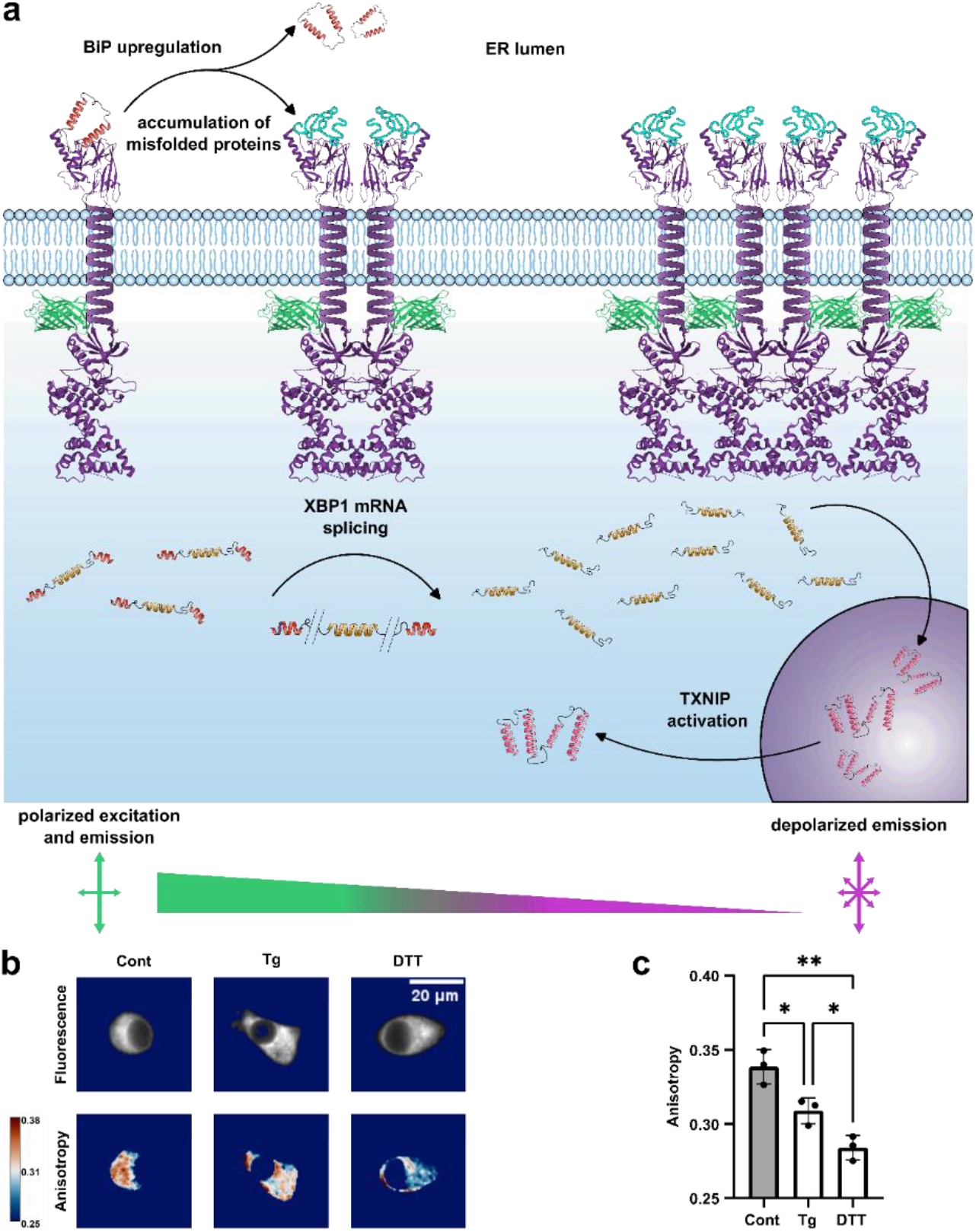
Apollo-IRE1 sensor design and validation of ER stress induced oligomerization. **(a)** Schematic representation of the Apollo-IRE1 homoFRET sensor and unfolded protein response (UPR). Top: Accumulation of misfolded proteins in the ER lumen triggers IRE1 activation, leading to three major downstream signaling outcomes: BiP upregulation, XBP1 mRNA splicing, and TXNIP activation. A cartoon showing the crystal structures of the IRE1 lumenal domain, kinase/RNase domain, and mVenus fluorescent protein connected by the transmembrane domain, which is represented only schematically as an amorphous section that connects all three crystal structures. The mVenus fluorescent protein is positioned at the cytoplasmic edge of the transmembrane domain. Bottom: The principle of homoFRET-based anisotropy sensing: monomeric sensor excited with polarized light emits polarized light (high anisotropy). Upon IRE1 dimerization, fluorescent proteins come within FRET distance (<10 nm), resulting in homoFRET and depolarization of emission (low anisotropy). **(b)** Representative fluorescence intensity (top row) and calculated anisotropy images (bottom row) of INS1E cells expressing Apollo-IRE1 under control conditions (Cont) or following treatment with thapsigargin (Tg; 1 µM, 6 h) or DTT (5 mM, 3 h). Anisotropy is displayed on a pseudocolor scale ranging from 0.25 to 0.38. Scale bar, 20 µm. **(c)** Quantification of Apollo-IRE1 anisotropy under control, Tg, and DTT treatment conditions. Data are presented as mean ± S.E.M., *n* = 3 independent experiments. \**P* < 0.05, \*\**P* < 0.01; statistical significance assessed by ordinary one-way ANOVA followed by Tukey’s multiple comparisons test (95% CI).

### Apollo-IRE1 Distinguishes Distinct Oligomeric States Under ER Stress

Our data and previous work suggest Tg and DTT induce distinct modes of ER stress with different oligomerization dynamics.^14,22^ To measure the aggregation state of Apollo-IRE1 in response to these chemical inducers of ER stress, we used progressive photobleaching combined with analysis of the impact on the apparent fluorescence anisotropy (i.e., the enhancement curves) according to established procedures^33,34^ (**Figure 2**). As fluorescent proteins photobleach, the shape of the enhancement curve depends on whether they form apparent monomers, dimers, or higher-order oligomers (**Figure 2a**). Non-FRETing monomeric fluorescent proteins show no change in fractional fluorescence with progressive photobleaching, so the anisotropy remains flat. In contrast, dimers and higher-order oligomers show linear responses with a slope and exponential responses, respectively (**Figure 2a**). Consistently, control cells showed a high Apollo-IRE1 fluorescence anisotropy that was relatively flat with progressive photobleaching, consistent with the sensor being primarily in a monomeric or non-FRETing resting dimeric state (**Figure 2b**). In contrast, cells treated with thapsigargin (Tg) showed a lower initial fluorescence anisotropy (i.e., at 1.0 fractional intensity) that increased linearly with progressive photobleaching (**Figure 2c**), whereas cells treated with DTT showed an exponential increase, consistent with high-order oligomers (**Figure 2d**). The linearity found in Tg-treated cells is consistent with the formation of homoFRETing dimers. In contrast, the DTT curve fit to an average oligomeric state of N=5 with low goodness-of-fit (*R*^2^ = 0.11), suggesting significant heterogeneity in the oligomeric population. Overall, these data are consistent with Tg-induced dimers and DTT-induced mixture of higher-order oligomers.

**Figure 2.**
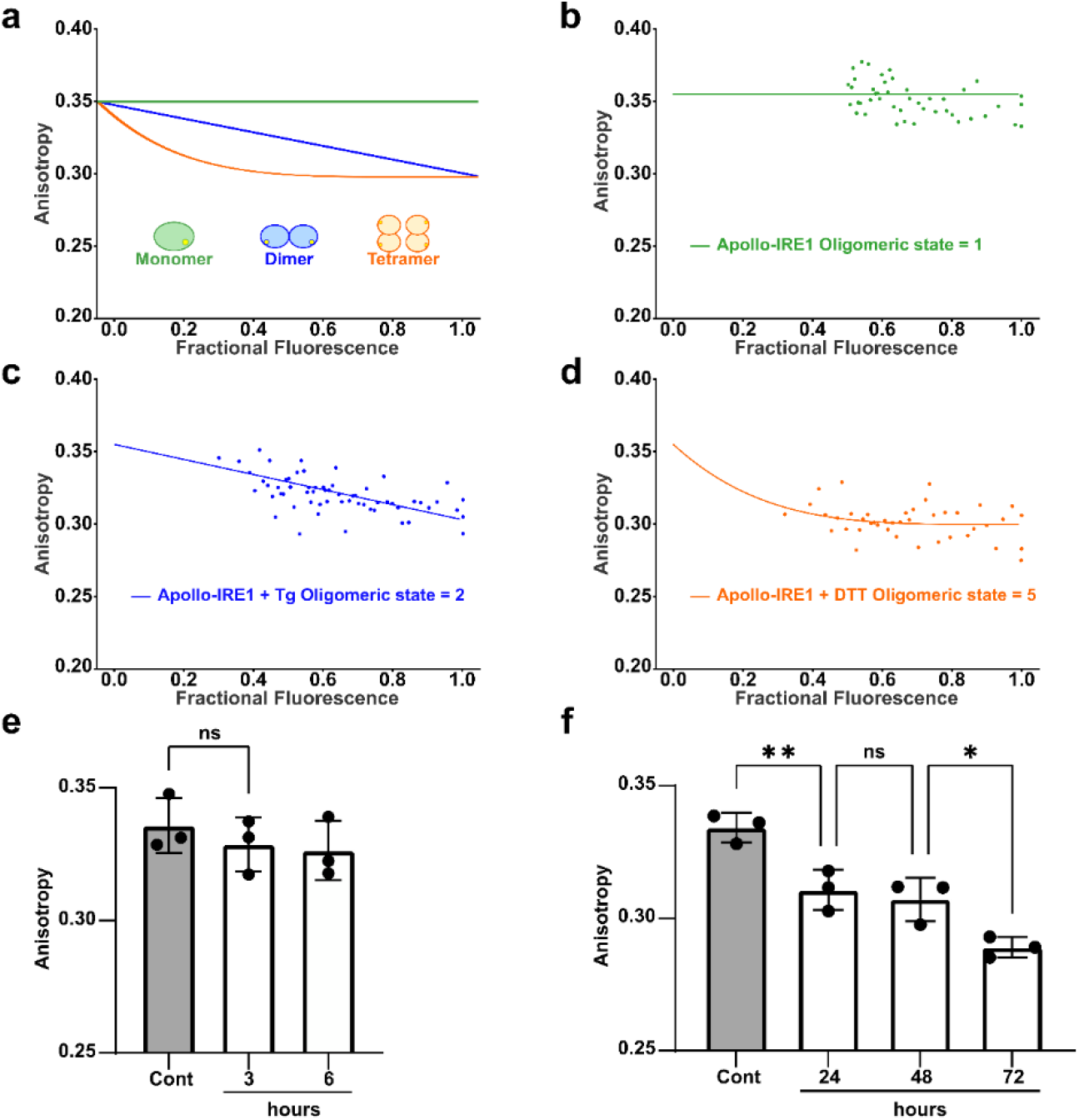
Validation of Apollo-IRE1 oligomeric state changes using enhancement curve analysis and physiologically induced ER stress response. **(a)** Schematic of the enhancement curve method for determining oligomeric state. Fractional fluorescence (x-axis) is calculated by normalizing observed fluorescence intensity to the expected signal for different oligomeric states. For monomeric proteins (oligomeric state = 1), anisotropy remains constant across all fractional fluorescence values (flat line). For dimeric or higher-order oligomers, anisotropy decreases with fractional fluorescence following a power-law relationship, where the curvature reflects the oligomeric state number, dimers show a near-linear decay while higher-order oligomers exhibit progressively steeper curvature at low fractional fluorescence values. Theoretical curves for monomer, dimer, and tetramer are illustrated. **(b)** Enhancement curve analysis of Apollo-IRE1 under control conditions showing a flat line consistent with predominantly monomeric sensor (oligomeric state = 1). **(c)** Enhancement curve analysis following thapsigargin treatment (1 µM, 6 h) showing a linear relationship consistent with dimeric IRE1 (oligomeric state = 2). **(d)** Enhancement curve analysis following DTT treatment (5 mM, 3 h) showing an apparent pentameric/higher-order assembly (oligomeric state = 5). The irregular stoichiometry and lower goodness-of-fit suggest heterogeneous oligomeric populations, possibly reflecting interactions with endogenous IRE1. **(e)** Apollo-IRE1 anisotropy in INS1E cells treated with high glucose (30 mM) for 3 and 6 h to assess acute ER stress responses. **(f)** Apollo-IRE1 anisotropy following prolonged high glucose treatment (30 mM) for 24, 48, and 72 h to simulate chronic, physiologically relevant ER stress. Data are presented as mean ± S.E.M., n = 3 independent experiments. ns, not significant; *P < 0.05, **P < 0.01; statistical significance assessed by ordinary one-way ANOVA followed by Tukey’s multiple comparisons test (95% CI).

To determine if the Apollo-IRE1 sensor was responsive to more physiologically relevant changes in ER stress, INS1E cells were treated with high-glucose (30 mM) for short (3 and 6 h) (**Figure 2e**) and long (24, 48, and 72 h) durations (**Figure 2f**). The sensor anisotropy remained consistently high during short-term treatment, consistent with previous studies showing that glucose-stimulated ER stress transitions from adaptive to potentially terminal after prolonged high-glucose exposure (typically >12-24 h).^12,22,35,36^ In contrast, the sensor showed a graded drop in Apollo-IRE1 anisotropy at 24, 48, and 72 h. These data suggest a transient plateau of Apollo-IRE1 dimerization is reached between 24 and 48 h, followed by a further decrease in anisotropy consistent with higher-order oligomers at 72 h. Taken together, these results indicate that Apollo-IRE1 is a live-cell sensor under both chemical and physiological conditions.

### Apollo-IRE1 and Multiplexing Imaging

A primary rationale for using Apollo sensors is their single-color design, which facilitates multicolor imaging. One potential application is measuring ER stress relative to other signals measured via immunofluorescence, such as the terminal ER stress marker thioredoxin-interacting protein (TXNIP).^9,21^ To co-image TXNIP immunofluorescence, we first confirmed Apollo-IRE1 responses are maintained after fixation (**Figure 3a**). TXNIP is regulated post-translationally, with inactive TXNIP retained in the nucleus and active TXNIP in the cytoplasm.^21,37^ To verify the activation of TXNIP by chronic high ER stress, we imaged the cytoplasmic-to-nuclear intensity of TXNIP immunofluorescence in INS1E cells treated with moderate (30 mM glucose, 24 h) and high (5 mM DTT, 3 h) ER stress (**Figure 3b**). Consistently, the cytoplasmic-to-nuclear intensity of TXNIP was relatively unaffected by glucose but increased significantly after DTT-induced ER stress. To compare Apollo-IRE1 and TXNIP responses more directly, we measured both responses simultaneously across many INS1E cells treated to induce moderate ER stress (30 mM glucose, 24 h) and varying times of high ER stress (5 mM DTT for 3, 4, and 5 h) (**Figure 3c-e**). These data in representative images (**Figure 3c**), histograms (**Figure 3d**), and scatter plots (**Figure 3e**) show that 30 mM glucose reduces Apollo-IRE1 steady-state anisotropy, consistent with dimerization. However, even after this relatively long treatment, TXNIP activation (i.e., cytoplasmic-to-nuclear intensity) remained low. DTT, in contrast, induced a much lower Apollo-IRE1 anisotropy consistent with higher-order oligomers. Notably, TXNIP cytoplasmic-to-nuclear intensity increased progressively at these time-points, correlating with oligomeric Apollo-IRE1 anisotropy (i.e., terminal UPR) (**Figure S2**). Overall, these data demonstrate that higher-order oligomerization of Apollo-IRE1 correlates with activation of late-stage TXNIP, enabling distinction between moderate (intermediate anisotropy, low TXNIP) and severe stress (lowest anisotropy, high TXNIP) associated with adaptive and terminal UPR activation, respectively.

**Figure 3.**
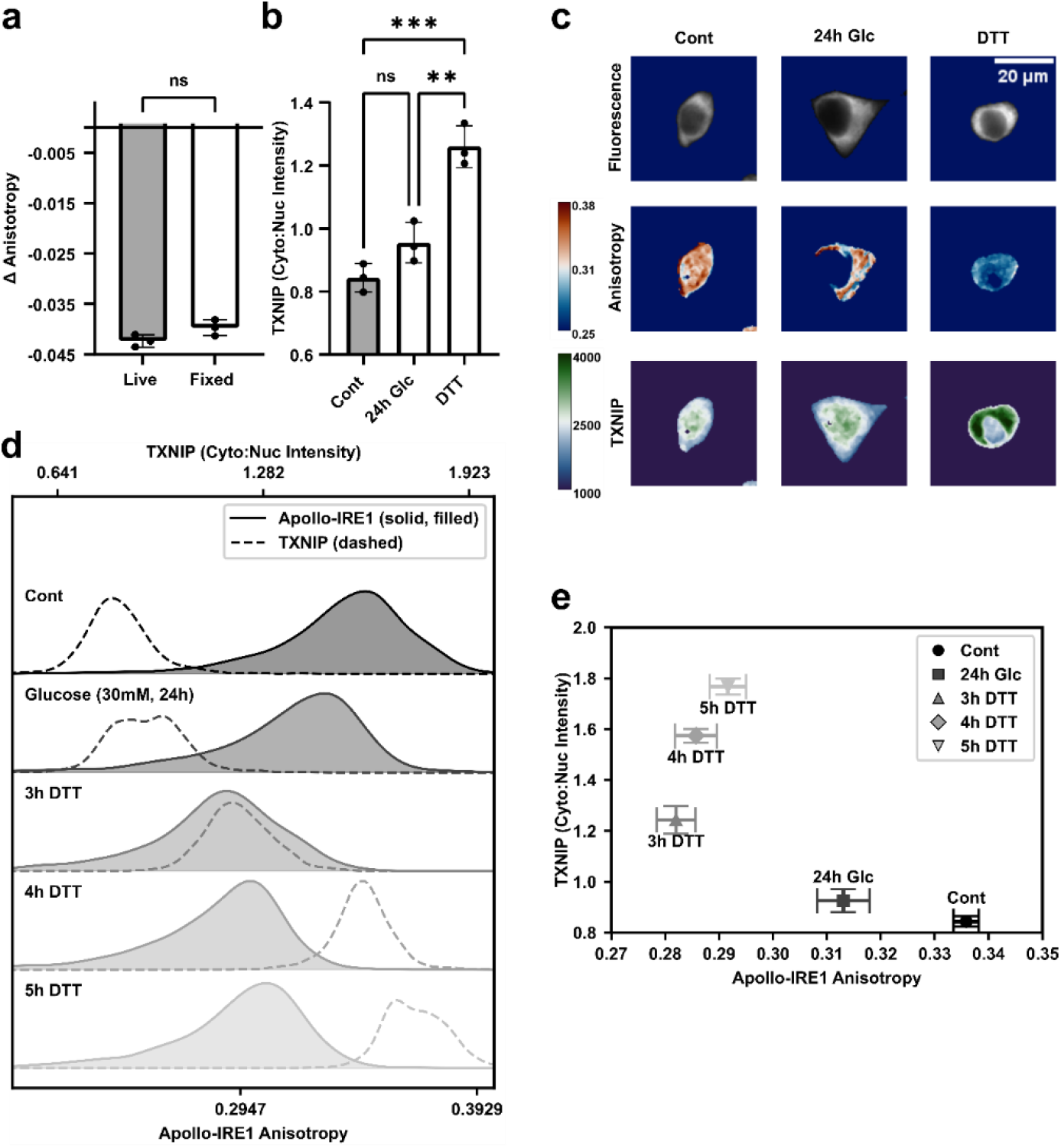
Apollo-IRE1 is compatible with paraformaldehyde fixation to enable multiplexed imaging with TXNIP immuno-fluorescence. **(a)** Comparison of Apollo-IRE1 anisotropy measured in live cells versus after paraformaldehyde fixation following DTT treatment (5 mM, 3 h). The change in anisotropy (Δ Anisotropy) relative to untreated controls is preserved after fixation, demonstrating compatibility with immunofluorescence workflows. ns, not significant; unpaired t-test (95% CI). **(b)** Quantification of TXNIP subcellular localization expressed as cytoplasmic-to-nuclear intensity ratio in INS1E cells under control conditions, following high glucose treatment (30 mM, 24 h), or DTT treatment (5 mM, 3 h). DTT treatment induces significant TXNIP nuclear-to-cytoplasmic translocation. ns, not significant; \*\**P* < 0.01, \*\*\**P* < 0.001; ordinary one-way ANOVA followed by Tukey’s multiple comparisons test (95% CI). **(c)** Representative multiplexed images of INS1E cells expressing Apollo-IRE1 fluorescence intensity (top row), Apollo-IRE1 anisotropy (middle row), and immunostained TXNIP intensity (bottom row) under control, high glucose (30 mM, 24 h), and DTT (5 mM, 3 h) conditions. Anisotropy is displayed on a pseudocolor scale (0.25–0.38). TXNIP fluorescence intensity is displayed on a pseudocolor scale (1000–4000 intensity units). Scale bar, 20 µm. **(d)** Ridgeline density plots showing the distribution of Apollo-IRE1 anisotropy (solid line, filled) and TXNIP cytoplasmic-to-nuclear intensity ratio (dashed line) across treatment conditions: control, high glucose (30 mM), and DTT time course (3, 4, and 5 h). Progressive DTT treatment shifts Apollo-IRE1 anisotropy slightly toward lower values (indicating increased oligomerization) while TXNIP cytoplasmic-to-nuclear ratio increases more dramatically. **(e)** Correlation between Apollo-IRE1 anisotropy and TXNIP cytoplasmic-to-nuclear intensity ratio across all treatment conditions. Each data point represents the mean of *n* = 3 biological replicates. Error bars represent S.E.M..

### Live-Cell Imaging of Apollo-IRE1

BiP (Binding Immunoglobulin Protein), also known as GRP78 (Glucose-Regulated Protein-78), is an ER-resident chaperone protein that both inhibits IRE1 oligomerization and assists in the proper folding of newly synthesized proteins.^14^ BiP inhibits IRE1α dimerization by binding to the protein’s lumenal domain, maintaining it in an inactive state. Under ER stress, BiP preferentially binds unfolded proteins, thereby allowing IRE1α dimerization. To investigate whether BiP levels competitively affect the kinetics of IRE1 activation, Apollo-IRE1-expressing INS1E cells were preincubated in low (2.5 mM) and high (30 mM) glucose to lower and raise BiP expression, respectively (**Figure S1b**), and then treated with DTT (**Figure 4a**). These data show that DTT induced a progressive decrease in anisotropy over 30 min, consistent with IRE1 activation, and that cells expressing higher levels of BiP showed a delayed response suggesting higher BiP expression delays IRE1α dimerization (**Figure 4b**). To confirm the previous response to DTT was related to BiP activity, we repeated the experiment using thapsigargin (Tg, 1 µM), which induces ER stress by depleting lumenal calcium, thereby impairing the calcium-dependent chaperone function of BiP.^38–40^ (**Figure 4c**). The Tg-treated cells showed a progressive decrease in anisotropy over 45 min, consistent with dimer formation, but, unlike DTT, did not display a significant difference between those with high and low BiP expression (**Figure 4c, d**). These data are consistent with BiP’s regulatory effect on IRE1α in the ER, and that calcium depletion overrides BiP-mediated regulation of Apollo-IRE1 activation. Together, these data suggest Apollo-IRE1 is a viable tool for measuring dynamic stress responses in living cells.

**Figure 4.**
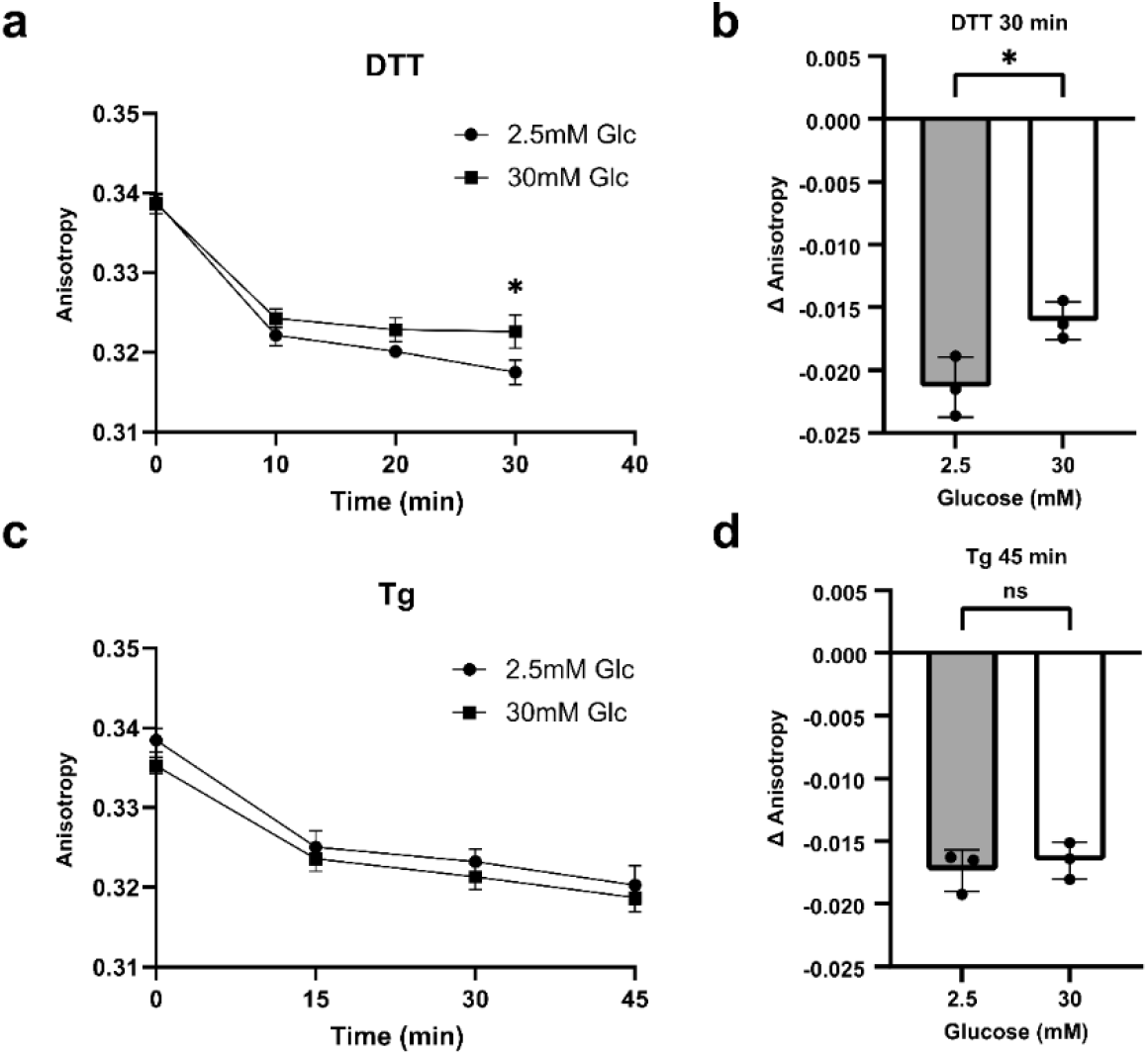
BiP expression levels modulate the kinetics of IRE1 oligomerization in response to ER stress. **(a)** INS1E cells were pre-incubated overnight in low (2.5 mM; low BiP) and high glucose (30 mM; high BiP) to lower and raise BiP levels, respectively, followed by imaging the Apollo-IRE1 anisotropy response to DTT treatment (5 mM). **(b)** Quantification of Apollo-IRE1 anisotropy change (Δ Anisotropy) at 30 min of DTT treatment, comparing low (2.5 mM) and high glucose (30 mM) pre-incubation. \**P* < 0.05; unpaired t-test (95% CI). **(c)** INS1E cells pre-incubated overnight in low (2.5 mM; low BiP) and high glucose (30 mM; high BiP) conditions followed by imaging the Apollo-IRE1 anisotropy response to thapsigargin (Tg; 1 µM). **(d)** Quantification of Apollo-IRE1 anisotropy change (Δ Anisotropy) at 45 min of thapsigargin treatment comparing low (2.5 mM) and high glucose (30 mM) preincubation. ns, not significant; unpaired t-test (95% CI). Data are presented as mean ± S.E.M., *n* = 3 independent experiments.

### Apollo-IRE1 Responses are Maintained in Primary Tissue

Immortalized beta-cell lines such as INS1E are valuable for establishing experimental models; however, they differ significantly from primary beta cells, including altered glucose sensitivity, insulin secretion dynamics, and stress response characteristics.^41–43^ To further evaluate the physiological relevance of Apollo-IRE1, we imaged responses in dispersed mouse islet cells transduced with adenoviruses expressing Apollo-IRE1 and mCherry under the rat insulin promoter (RIP). (**Figure 5**). The mCherry expression allowed us to quantify the Apollo-IRE1 responses only in insulin-expressing beta cells (**Figure 5a**). The Apollo-IRE1 sensor was expressed in <60% of cells, excluded from the nucleus, and exhibited a web-like ER localization throughout the cytoplasm, similar to that observed in INS1E cells. The beta cells showed a graded response to each ER stressor, both visually (**Figure 5a**) and in aggregate (**Figure 5b**). These data demonstrate that Apollo-IRE1 maintains its spatial and quantitative performance in a primary tissue model. Together, these results show the ability to dynamically measure physiological ER stress in tissue-derived single cells, demonstrating that Apollo-IRE1 is a valuable tool for investigating beta-cell dysfunction in diabetes models.

**Figure 5.**
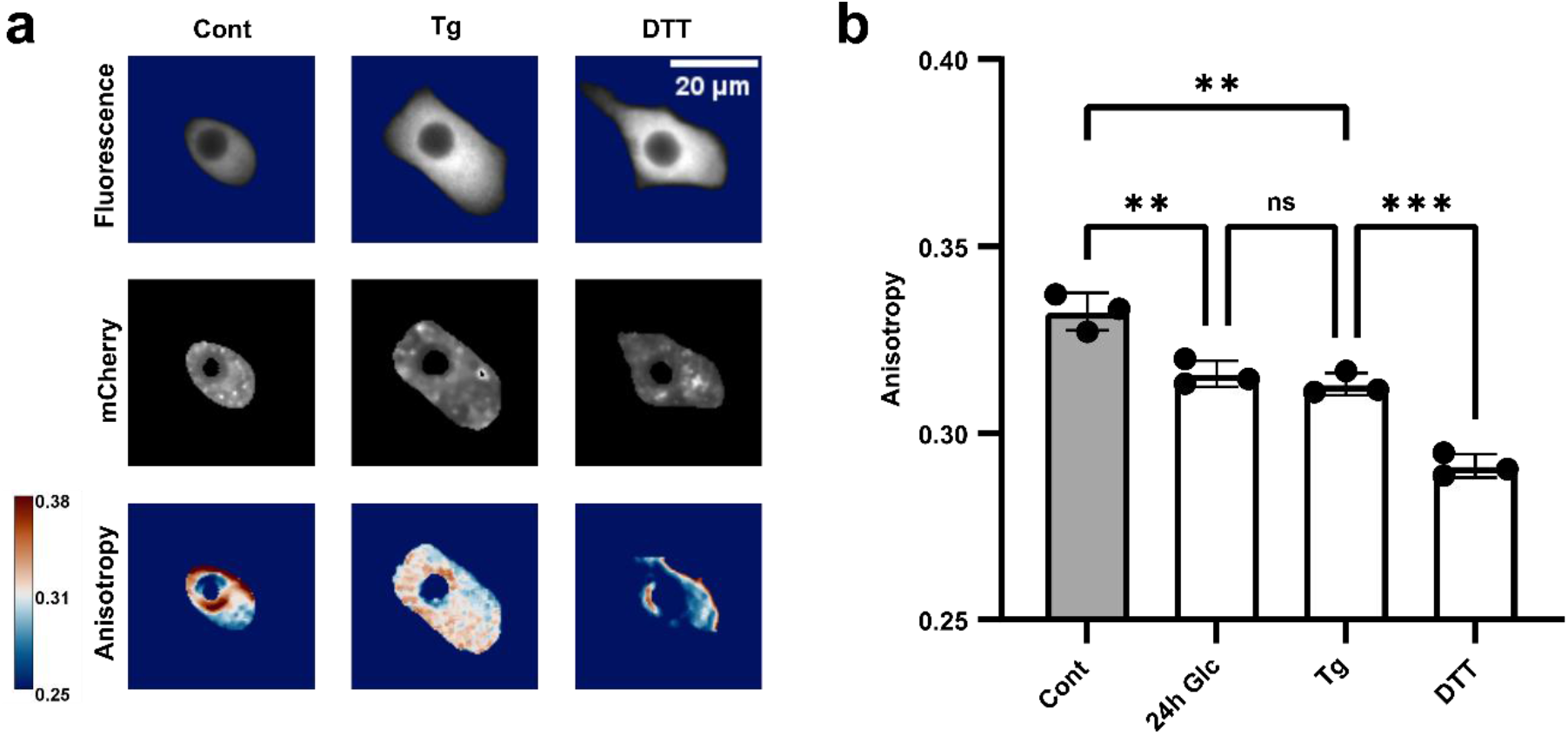
Apollo-IRE1 sensor validation in dispersed primary mouse islet cells. **(a)** Representative images of mouse islets (FVB/Nj, male) dispersed to single cells and transduced with Apollo-IRE1 and a rat insulin promoter (RIP)-mCherry reporter to identify beta cells. Columns show cells treated with control, thapsigargin (1 µM, 6 h), and DTT (5 mM, 3 h). Rows show Apollo-IRE1 fluorescence intensity (top), RIP-mCherry fluorescence (middle), and calculated Apollo-IRE1 anisotropy (bottom). Anisotropy is displayed using a pseudocolor scale (0.25–0.38). Scale bar, 20 µm. **(b)** Quantification of Apollo-IRE1 anisotropy in mCherry-positive beta cells treated with control, high glucose (30 mM, 48 h), thapsigargin (1 µM, 6 h), and DTT (5 mM, 3 h). Data are presented as mean ± S.E.M., *n* = 3 independent experiments. ns, not significant; \**P* < 0.05, \*\**P* < 0.01, \*\*\**P* < 0.001; statistical significance assessed by ordinary one-way ANOVA followed by Tukey’s multiple comparisons test (95% CI).

## DISCUSSION

Here, we developed Apollo-IRE1, a genetically encoded homoFRET sensor for live-cell imaging of ER stress based on the IF2L mutant of IRE1 from the Walter lab.^24^ This single-color sensor reports on ER stress-induced changes in IRE1 oligomerization state, with a narrow spectral bandwidth and ratiometric output. Our photobleaching and enhancement curve analysis revealed that Apollo-IRE1 distinguishes at least three apparent states: a baseline state in unstressed cells, a dimeric state under moderate stress, and higher-order oligomers under severe stress. The Apollo-IRE1 sensor response was also fixable. This, in combination with the single-color design, enabled multiplexed immunofluorescence imaging, which we used to correlate higher-order Apollo-IRE1 assembly with the terminal UPR marker TXNIP. The ratiometric readout of Apollo-IRE1 provided consistent measurements across multiple days and instrumentation setups. Finally, Apollo-IRE1 responses were maintained in primary dispersed mouse islet beta cells, demonstrating translational potential beyond immortalized cell lines. Together, these results establish Apollo-IRE1 as a practical tool for investigating the dynamic progression of ER stress in living cells, with relevance to understanding beta-cell dysfunction in diabetes.

Enhancement curve analysis revealed that Apollo-IRE1 distinguishes at least three apparent states: (i) monomers in unstressed cells, (ii) dimers under moderate stress (e.g., Tg and 24 h of high glucose), and (iii) higher-order oligomers under severe stress (e.g., DTT and 72 h of high glucose). Recent work suggests that endogenous inactive IRE1 exists as constitutive dimers,^23^ raising the question of why Apollo-IRE1 appears monomeric in unstressed cells. We postulate that the monomeric state results either from our use of the IF2L mutation, which may yield inactive true monomers, or from the formation of inactive dimers that conformationally place the fluorescent proteins outside the Förster distance. Activation in the latter model would necessitate a conformational change that subsequently results in homoFRET. It should be noted that the precise conformational changes within dimers upon activation have not been directly measured using the intrasequence fluorescent protein configuration employed in Apollo-IRE1. Regardless of the precise molecular mechanism, Apollo-IRE1 reliably distinguishes stress states, as evidenced by rapid responses to stressors and by its correlation with TXNIP. Our BiP dynamics experiments provide further mechanistic insight: high BiP expression delayed the kinetics of the Apollo-IRE1 response to DTT, whereas low BiP permitted a faster response. These data are consistent with the direct ligand model of IRE1 activation, in which unfolded proteins bind directly to IRE1’s lumenal domain, while BiP competes for binding and sets the sensitivity threshold. Notably, thapsigargin bypassed modulation by BiP entirely, suggesting that depletion of calcium in the ER may directly release BiP from IRE1 or destabilize the inactive conformation through another mechanism. These findings position Apollo-IRE1 as a sensor of both adaptive- and terminal-UPR and suggest that BiP acts as a modulator of sensitivity rather than the primary activation trigger.

The Apollo-IRE1 sensor surprisingly formed higher-order oligomers under severe stress despite having the IF2L mutation. This mutation disrupts the IF2L lumenal domain interface, preventing the formation of foci, while preserving the IF1L lumenal domain interface, which is necessary for dimerization. We therefore anticipated the sensor would not oligomerize. However, our data showing oligomerization could indicate a lower level of oligomerization (i.e., below the threshold for forming foci) driven by a different portion of IRE1. The trans-membrane domain of IRE1α has been shown to form oligomers independently in membrane-mimetic environments.^44^ Since the IF2L mutation does not affect this domain, it may contribute to the observed higher-order assemblies. Additionally, we postulate that endogenous IRE1 could be influencing sensor readout under severe stress conditions. First, DTT treatment produced an enhancement curve best fit by a pentameric stoichiometry, inconsistent with the canonical dimer-stacking model of IRE1 oligomerization, which predicts even-numbered oligomers. This irregular stoichiometry suggests that oligomerizing endogenous IRE1 bridges multiple Apollo-IRE1 molecules into FRET range through heterodimerization. Second, the lower goodness-of-fit for the DTT condition compared to thapsigargin suggests greater heterogeneity in the oligomeric population under severe stress. This effect may be particularly pronounced in secretory cells such as INS1E beta cells, which express high levels of endogenous IRE1 to meet the demands of insulin production.

Apollo sensor design offers several key advantages. First is the ratiometric, intensity-independent readout. Fluorescence anisotropy is calculated from the ratio of parallel and perpendicular emission, making it insensitive to variations in expression level, illumination power, and instrumentation differences. This enabled consistent baseline measurements across multiple experimental days with low variability, enabling longitudinal studies and high-content imaging. Second, the single-color design leaves substantial spectral bandwidth for multiplexing, unlike heteroFRET sensors that consume much of the visible spectrum. We demonstrated this by simultaneously imaging Apollo-IRE1 with TXNIP immunofluorescence and with mCherry in dispersed islets. Third, the Apollo-IRE1 sensor could be fixed. This enabled endpoint immunofluorescence for markers that cannot be imaged in real time. These practical features position Apollo-IRE1 to offer quantitative, continuous readout of early activation states with spectral flexibility for multiparametric imaging. Finally, like all live-cell sensors, Apollo-IRE1 enables the capture of dynamic changes on time-scales of minutes to hours that are inaccessible to destructive methods such as western blotting, while single-cell resolution reveals heterogeneity masked by population averaging. The slow dynamics of ER stress progression, combined with anisotropy stability, allow samples to be removed and returned to the microscope without losing baseline reference.

We envision several future applications for the Apollo-IRE1 sensor. Notably, the single-color design enables co-expression with other genetically encoded sensors. For instance, co-expression with Apollo-NADP^+^ could reveal the temporal relationship between NADPH/NADP^+^ redox state and ER stress resistance, as the thioredoxin antioxidant system requires NADPH to manage oxidative stress during protein folding.^45^ Since TXNIP inhibits thioredoxin and links terminal UPR to inflammasome activation, simultaneous imaging could capture this regulatory relationship in real time.^13,21^ Our BiP dynamics data suggest that chaperone levels tune the sensitivity of IRE1 activation. Thus, future experiments could co-image Apollo-IRE1 with fluorescently tagged BiP to assess competitive binding dynamics under different stress conditions.^14,46^ The calcium dependence observed with thapsigargin represents another avenue for investigation, given that ER calcium dysregulation is implicated in beta-cell dysfunction during diabetes progression. Apollo-IRE1 maintains its response in dispersed primary mouse islet beta cells, and the Apollo sensor family has been validated in vivo using zebrafish models where Apollo-NADP^+^ provided stable anisotropy measurements across tissue depth.^30^ These precedents suggest Apollo-IRE1 could be adapted for in vivo imaging of ER stress in intact islets or transplanted beta cells, providing direct sensing of early IRE1 activation events rather than relying on downstream markers.

### LIMITATIONS

#### Our work has several limitations

(1) Enhancement curves report population-level distributions rather than single-cell oligomeric states. Single-cell enhancement curves exhibit higher variance and cannot conclusively determine the oligomeric state of an individual cell. Enhancement curve fitting approximates the distribution of states across a population, and even under severe stress, the majority of Apollo-IRE1 molecules remain in the baseline state, indicating the sensor detects shifts in a subset of molecules rather than wholesale conversion. (2) Apollo-IRE1 dynamic range may be limited by the inherent constraints of fluorescence anisotropy measurements and hetero-dimerization with endogenous IRE1. Anisotropy values are bounded by theoretical limits, yielding a narrower dynamic range than intensity-based measurements. Heterodimerization further compresses this range by forming single-fluorophore dimers that appear as monomers and mixed oligomeric populations with suboptimal fluorophore geometry for efficient homoFRET, an effect particularly relevant in pancreatic beta cells, which express high levels of endogenous IRE1. Dynamic range may be improved by using cell lines with low endogenous IRE1 expression or increasing Apollo-IRE1 expression to favor sensor homodimers. And finally, (3) the IF2L mutant was shown by Karagöz et al.^24^ to have no XBP1 splicing or kinase activity when expressed in IRE1-knockout cells; this was not independently verified in our hands.

## MATERIALS AND METHODS

### Cell Culture, Transfection, and Treatments

INS1E beta cells were cultured in RPMI-1640 medium supplemented with 5% fetal bovine serum (FBS), 1 mM sodium pyruvate, 10 mM HEPES, 50 µM 2-mercaptoethanol, penicillin (100 U/mL), and streptomycin (100 U/mL), under humidified 5% CO_2_ at 37°C. For imaging, cells were plated on no. 1.5 glass-bottom dishes (MatTek Corporation, Ashland, MA). The following day the cells were either transfected with Apollo-IRE1 plasmid (0.5–1.0 µg/dish/plasmid) using Lipofectamine 3000 following the manufacturer’s protocol or transduced with Apollo-IRE1 adenovirus (2 × 10^7^ IFU/mL) (VectorBuilder, ID: VB210813-1003dnf) and incubated for 24 h. After incubation, the media was replaced with fresh culture media, and the cells were left to recover for a further 24 h prior to treatment and/or imaging. Chemical treatments were introduced into the culture medium and imaging buffer at the specified time points.

### Mouse Islet Isolation, Dispersion, and Transduction

This animal procedure received approval from the Animal Care Committee of the University Health Network (Toronto, ON, Canada), adhering to guidelines established by the Canadian Council on Animal Care (Animal Use Protocol #1531). Pancreatic islets were surgically extracted from 8–12 week-old male FVB mice (Jackson Laboratory) and subjected to collagenase digestion (Roche Applied Science). After digestion, islets were individually selected by pipetting under a microscope and placed into a 1.5 mL Eppendorf tube containing 50 µL of RPMI islet media (10% FBS) and 50 µL trypsin/ethylenediaminetetraacetic acid (EDTA). These islets were then placed in a 37°C water bath and manually agitated for 10–15 min. The resulting dissociated islet mixture was diluted to a final volume of 1 mL using RPMI islet media and pipetted into glass-bottom plates. For cell adhesion, the glass-bottom plates (24 or 48 well) were pre-coated with poly-D-lysine hydrobromide (Sigma-Aldrich, #P6407) and placed in an incubator for 1 h at 37°C. After plating, cells were transduced with adenovirus vector (1:10 dilution or 2 × 10^7^ IFU/mL) and incubated for 24 h. After incubation, the media was replaced with fresh culture media, and the cells were left to recover for a further 24 h before treatment and/or imaging.

### Molecular Cloning of IRE1 Constructs

A mutant version of IRE1 was previously created, human IRE1α-IF2-GFP, with a mutated oligomerization site (WLLI359-362 to GSGS359-362; ‘IF2L mutant’) to only allow IRE1 dimerization.^24^ A homoFRET modification of human IRE1α-IF2-GFP, coined Apollo-IRE1, was cloned into a pcDNA3.1(+) backbone with mVenus and mCerulean replacing the GFP (**Figure S3**). The FP was placed intrasequence in the IRE1 construct. First, the lumenal section was digested using restriction enzymes EcoRI and HindIII and inserted into an empty pcDNA3.1(+) backbone. The IRE1 kinase and RNAse domain was next cut out using NotI and XbaI restriction enzymes and homologous recombination was used to combine the lumenal and kinase/RNAse sections. The new pcDNA-IRE1 construct was cut using NotI and EcoRI restriction enzymes and homologous recombined with mVenus and mCerulean. The following primers were used throughout the cloning process: IRE1-EcoRI, 5’-ATGCCGGCCCGGCGGCTGCT-3’; IRE1-HindIII, 5’-GGGAGGGTCTGAGGAAGGTG-3’; IRE1-NotI, 5’-CATGCATCAGCAGCAGC-3’; IRE1-XbaI, 5’-GCACACGGCAAGATCAAGGC-3’. The final constructs were confirmed with Sanger sequencing. Two final versions of Apollo-IRE1 were produced: mVenus-tagged Apollo-IRE1 and mCeru-lean-tagged Apollo-IRE1.

### XBP1 mRNA Splicing

Total RNA was isolated from INS1E cells using TRIzol reagent (Invitrogen). The resulting RNA was reverse transcribed to cDNA by the High-Capacity cDNA Reverse Transcription Kit (Applied Biosystems). Primers designed to flank the XBP1 intron excised by IRE1 exonuclease activity were 5’-AAA CAG AGT AGC AGC ACA GAC TGC-3’ and 5’-TCC TTC TGG GTA GAC CTC TGG GAG-3’ (IDT DNA). The cDNA was amplified by RT-PCR (Qiagen OneStep RT-PCR Kit). Amplification thermal cycling conditions were: 50°C (30 min), 95°C (15 min), 30 cycles of 94°C (1 min), 62°C (1 min), 72°C (1 min), 72°C (10 min). Amplified DNA fragments were separated by using 3% agarose gel by electrophoresis and stained with ethidium bromide. Spliced form of XBP1 appears lower on the gel since uXBP1 is 480 bp and sXBP1 is 454 bp.

### TXNIP Immunofluorescence

Following treatment with either DTT or high glucose (30 mM), cells were washed with phosphate-buffered saline (PBS; Gibco) and fixed with 2% paraformaldehyde solution (prepared from 95% PFA powder, Sigma Aldrich) in PBS (pH 7.4) for approximately 20 min at room temperature. After fixation, cells were permeabilized using an antibody dilution (AD) buffer consisting of 0.1% Triton X-100 (Sigma Aldrich) in PBS (prepared by diluting 50 µL of Triton X-100 in 50 mL PBS). Next, a blocking solution was prepared containing 0.2% bovine serum albumin (BSA; Bishop) and 5% normal goat serum (NGS; Cell Signaling Technologies) in PBS. BSA solution was prepared separately at a concentration of 1 mg/mL by dissolving 40 mg of BSA in 40 mL of PBS. Permeabilization was performed by aspirating the fixative and adding 1 mL of the AD buffer for 5 min. The AD solution was then aspirated and replaced with 200 µL of blocking solution per dish. Cells were incubated with the blocking solution for 1 h at room temperature in the dark (covered in aluminum foil).

Primary antibody solution was prepared using 0.2% BSA (4 µL), 1.5% NGS, and 1:100 dilution of a recombinant monoclonal TXNIP primary antibody (ThermoFisher, Cat# MA5-32771) in AD buffer. Following the blocking step, cells were washed once with 1 mL PBS, which was then aspirated and replaced with 200 µL of the primary antibody solution. Incubation with the primary antibody was performed for 1 h at room temperature in the dark. Secondary antibody solution was prepared by mixing 30 µL of NGS (1.5%), 4 µL of 0.2% BSA solution, and 4 µL of goat anti-rabbit Alexa Fluor 568 secondary antibody (Invitrogen) at a 1:500 dilution, in AD buffer. After primary antibody incubation, cells were washed eight times with PBS, and then 200 µL of the secondary antibody solution was added to each dish. Secondary antibody incubation was carried out for 1 h at room temperature in the dark. Following secondary antibody incubation, cells were washed three times with PBS. Imaging was performed either immediately by adding 2 mL of PBS (or imaging buffer) before transferring dishes to the stage-top microscope, or the stained samples were stored overnight at 4°C in a humidified chamber wrapped in aluminum foil for imaging the next day.

### GRP78/BiP Western Blot

All immunoblots were performed and analyzed as described in Ting et al.^32^ In brief, after incubation in 2.5, 5, 11, and 30 mM glucose, INS1E cells were lysed in ice-cold RIPA buffer (1% NP40, 0.1% SDS, 0.5% deoxycholate in PBS, supplemented with 1X complete™, EDTA-free Mini Protease Inhibitor Cocktail (Sigma Cat#11873580001)) for 15 min. Protein concentrations in lysates were determined by Protein Assay Dye Reagent (BioRad #5000006), diluted in 2 × Laemmli sample buffer (BioRad Cat#161–0737) with fresh 2-mercaptoethanol (BioRad #1610710), and heated at 95°C for 5 min. Samples (20 µg of protein per lane) were resolved on 8% SDS-PAGE gels and transferred to polyvinylidene difluoride membranes (Sigma #IPVH00010) using a wet transfer system. Membranes were blocked with 3% BSA (Bioshop #ALB003) in Tris-buffered saline-Tween (TBST) for 1 h at room temperature. Membranes were incubated with primary antibodies overnight: anti-GRP78/BiP (Sigma, G8918) and anti-Actin (Sigma, A2066), followed by washing and incubation with HRP-conjugated anti-rabbit IgG (CST#7074) (22°C, 1 h). Blots were developed using Immobilon Forte Western HRP substrate (Sigma, WBLUF0100), imaged with Microchemi 4.2 (BioRad) and analyzed with ImageJ.

### Microscope Set-Up

Images were collected using a custom-built wide-field RAMM microscope (ASI) equipped with three excitation LEDs (405, 505, and 590 nm), an excitation polarizing filter (Edmund Optics, Barrington, NJ), and a 40×/0.75 NA air objective lens (Olympus, Richmond Hill, Canada). Fluorescence was directed through a Venus (ET535/30m), mCherry (ET632/60m), or Cerulean (ET470/24m) emission filter mounted on a filter wheel and then split into parallel (I_||_) and perpendicular (I_⊥_) intensity using an Optosplit II (Cairn, Faversham, UK). This setup allowed for the simultaneous collection of I_||_ and I_⊥_ on distinct regions of an IRIS 15 CMOS camera (Teledyne Photometrics, Tucson, AZ). To reduce photobleaching, images were acquired using 4×4 binning, 80% LED power, and moderate integration time (800–1200 ms). The culture media was replaced 1 h prior to imaging with BMHH imaging buffer (125 mM NaCl, 5.7 mM KCl, 2.5 mM CaCl_2_, 1.2 mM MgCl_2_, 0.1% BSA, and 10 mM HEPES at pH 7.4). The microscope stage top incubator (Okolab) was controlled for humidity and temperature (37°C).

### Image Analysis

Steady-state fluorescence anisotropy images were analyzed as previously described.^47^ All images were analyzed using Fiji-ImageJ software^48^ and custom plugins, available at https://github.com/Rocheleau-Lab/Optosplit-Anisotropy-Analysis-scripts. Anisotropy (r) was calculated pixel-by-pixel using images background corrected by a rolling ball filter with radius set to approximately the size of a single cell in an image (100 pixels). Anisotropy was calculated as r = (I_||_−*G*I_⊥_)/(I_||_+2*G*I_⊥_). The G-factor was measured using a standard fluorescein solution with a negligible anisotropy (5 mM dissolved in PBS solution), thus allowing *G* to be calculated from fluorescein images as *G* = I_||_ / I_⊥_. For images collected using the 1.42 NA objective lens, additional correction factors (Ka, Kb, and Kc) were implemented to account for mixing of polarized light as a result of wider detection angles. ROIs were drawn either manually or by using an implementation of CellPose algorithm to encircle whole cells.^49^ Additionally, cell segmentation was achieved using Fiji-ImageJ software (**Figure S4**).

### Oligomerization Enhancement Curves

To measure the degree of oligomerization of Apollo-IRE1, enhancement curves were collected and analyzed based on previously described methods.^34^ Briefly, 2P (950 nm excitation) fluorescence anisotropy images were collected, followed by 10 frames of sequential photobleaching using the one-photon laser (514 nm at 30-40% power). This cycle of anisotropy imaging and photobleaching was repeated until cells were 60% photobleached (∼10 cycles). The anisotropy for cellular regions of interest were plotted against the photobleached fluorescence intensity (I/Io where I is the current fluorescence intensity and Io is the initial fluorescence intensity). The curves were fit to the following equation:^34^

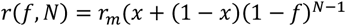

Where r_m_ is the anisotropy value of a monomeric fluorophore, x is the fraction of fluorophore in a monomeric state, N is the minimum oligomerization state, (1-x) is the fraction of fluorophore in oligomeric state N, f is the fractional fluorescence, and r is the resulting anisotropy. Values for N and x were fit using Python and scipy.optimize.curve_fit function.

### Anisotropy Cell Images

Anisotropy inherently has a low dynamic range making it difficult to depict results visually. Images are made meaningful through use of proper color mapping and thresholding. For anisotropy images presented throughout this paper the following steps were taken using ImageJ software. First the images were cropped, and background was removed using automatic thresholding (Otsu Dark) of the parallel intensity image. A median filter with radius 1 was applied to clean the images. Next, the lookup table was set to Vik (Fabio Crameri) and set to min/max values of 0.25/0.38. A script was developed to complete this process which can be found at (https://github.com/Rocheleau-Lab).

### Statistical Analysis

All data are presented as mean ± standard error of the mean (S.E.M.) where the number of independent experimental repeats is at least 3 (*n* ≥ 3). Each independent experiment consists of at least 30 cells (30 technical replicates). Anisotropy measurements were confirmed as normally distributed using a normal probability plot. Statistical significance was determined using Prism6 (GraphPad, SanDiego, CA). In comparing three or more groups, statistical significance was determined using one-way analysis of variance (ANOVA) followed by a Tukey’s multiple comparisons post-hoc test.

## ASSOCIATED CONTENT

### Supporting Information

This submission includes Supporting Information containing four supplemental figures (Figures S1–S4) with additional validation data and methodological details.

## AUTHOR INFORMATION

### Present Addresses

**Eric J. Floro** − Institute of Biomedical Engineering, University of Toronto, Toronto, Ontario M5S 3G9, Canada; Toronto General Hospital Research Institute, University Health Network, Toronto, Ontario M5G 2C4, Canada;

**Alex M. Bennett** − Institute of Biomedical Engineering, University of Toronto, Toronto, Ontario M5S 3G9, Canada; Toronto General Hospital Research Institute, University Health Network, Toronto, Ontario M5G 2C4, Canada

**Romario Regeenes** − Institute of Biomedical Engineering, University of Toronto, Toronto, Ontario M5S 3G9, Canada; Toronto General Hospital Research Institute, University Health Network, Toronto, Ontario M5G 2C4, Canada;

**Huntley H. Chang** − Institute of Biomedical Engineering, University of Toronto, Toronto, Ontario M5S 3G9, Canada; Toronto General Hospital Research Institute, University Health Network, Toronto, Ontario M5G 2C4, Canada;

**Nitya Gulati** − Institute of Biomedical Engineering, University of Toronto, Toronto, Ontario M5S 3G9, Canada; Toronto General Hospital Research Institute, University Health Network, Toronto, Ontario M5G 2C4, Canada;

### Notes

The authors declare no competing financial interest. The authors declare no competing interests.

## FOR TABLE OF CONTENTS ONLY

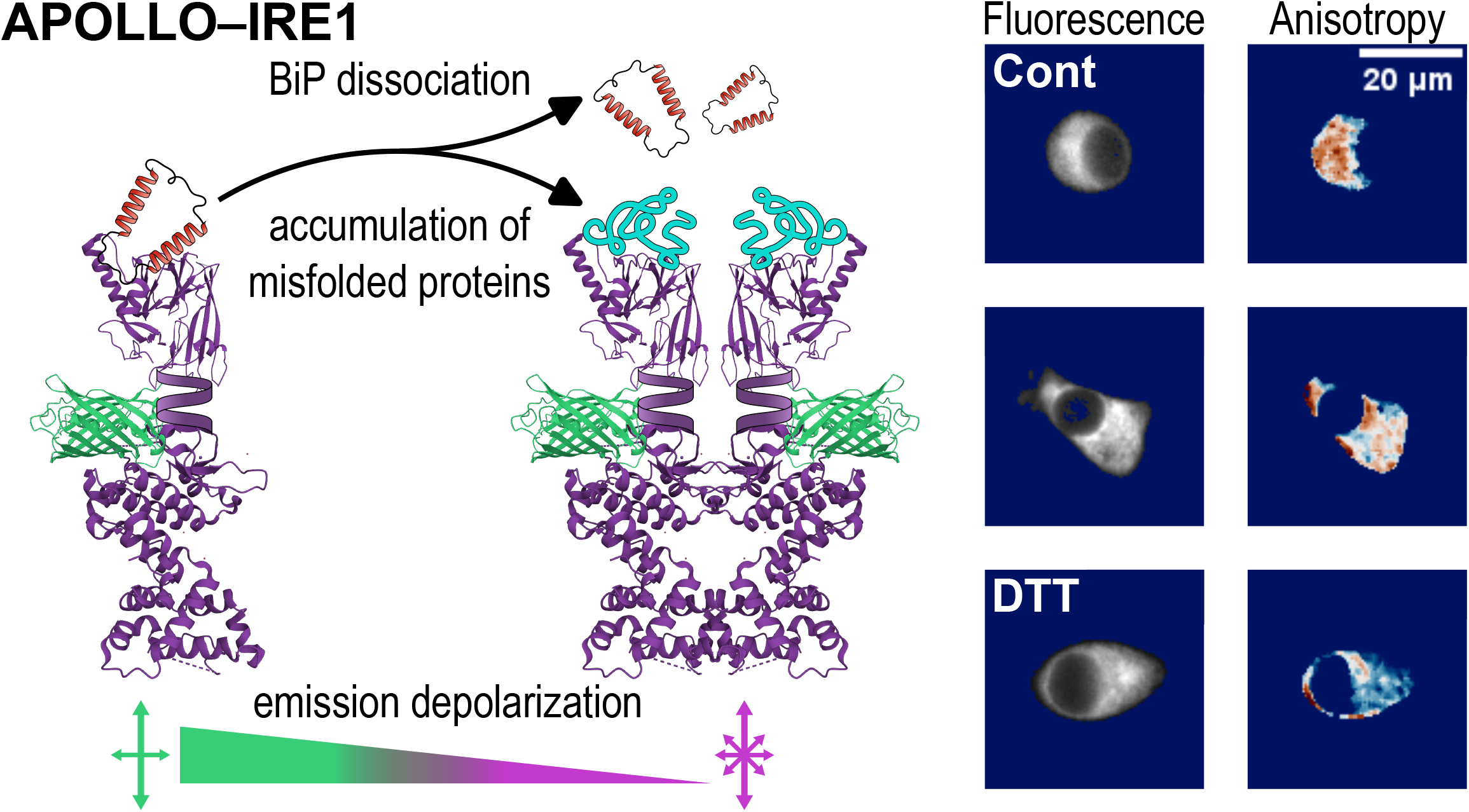

