## Supplemental Information for "Apollo-IRE1: A Genetically Encoded Sensor for Live Cell and Multiplexed Imaging of ER Stress"

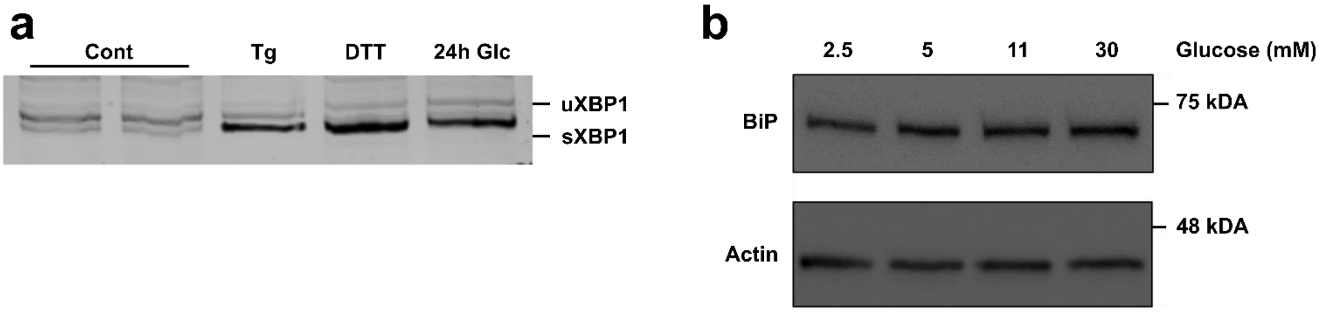


**Figure S1. Validation of IRE1 activation by XBP1 splicing analysis. (a)** RT-PCR analysis of XBP1 mRNA splicing in Apollo-IRE1 expressing INS1E cells. Primers flanking the 26-nucleotide intron excised by activated IRE1 endonuclease activity distinguish unspliced XBP1 (uXBP1; upper band) from spliced XBP1 (sXBP1; lower band). Lanes show: control (Cont), thapsigargin treated (Tg; 1 μM, 6 h), DTT-treated (5 mM, 3 h), and high glucose-treated (30 mM, 24 h) conditions. DTT treatment induces robust XBP1 splicing, confirming IRE1 activation. High glucose and Tg treatments show modest XBP1 splicing consistent with mild ER stress induction. **(b)** Western blot analysis of BiP protein expression in INS1E cells following overnight incubation in varying glucose concentrations (2.5, 5, 11, and 30 mM). Actin serves as loading control. BiP expression increases with glucose concentration, consistent with glucose-dependent modulation of ER chaperone capacity. Molecular weight markers are indicated (BiP: 75 kDa; Actin: 48 kDa).

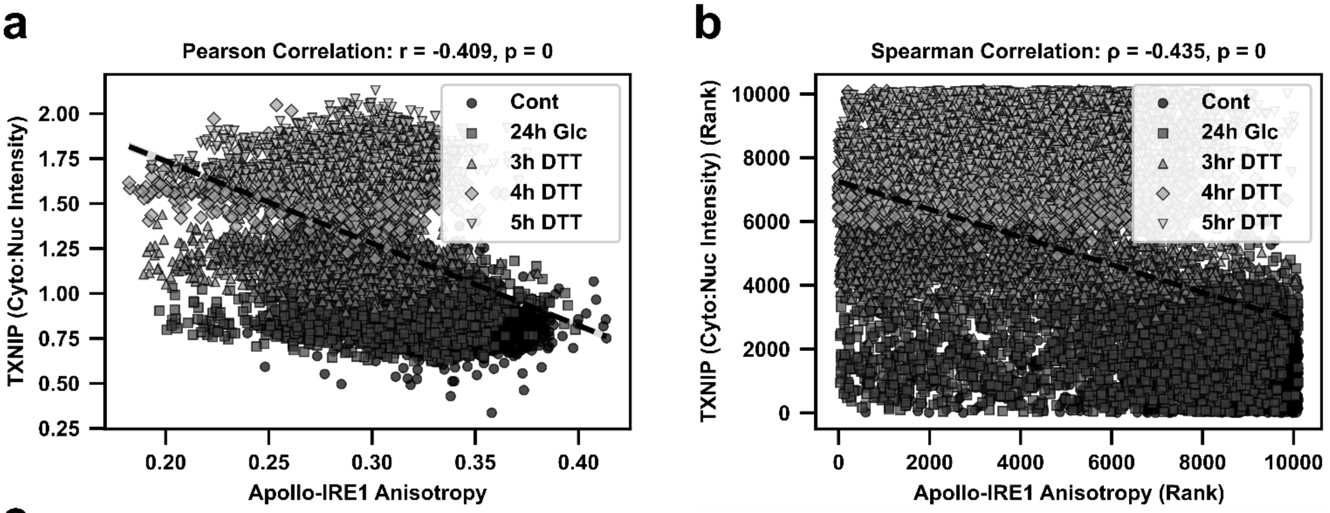


**c**

**Summary of Correlation analysis Between TXNIP (Cyto:Nuc Intensity) and Apollo-IRE1 Anisotropy**

| Correlation Type | Data Level | Sample Size (N) | Correlation Coefficient | p-value | Interpretation |
|---|---|---|---|---|---|
| Pearson | Per-replicate means | 15 | r = -0.741 | 0.0016 | Strong negative linear correlation across biological replicates |
| Pearson | Raw measurements | > 10,000 | r = -0.409 | < 0.0001 | Moderate negative linear correlation across all measurements |
| Spearman | Per-replicate means | 15 | ρ = -0.646 | 0.0092 | Strong negative monotonic correlation across replicates |
| Spearman | Raw measurements | > 10,000 | ρ = -0.435 | < 0.0001 | Moderate negative monotonic correlation across all measurements |

**Figure S2. Statistical correlation analysis between Apollo-IRE1 anisotropy and TXNIP fluorescence. (a)** Pearson correlation analysis of TXNIP cytoplasmic-to-nuclear intensity ratio versus Apollo-IRE1 anisotropy. Per-replicate means ($n$ = 15; 3 biological replicates × 5 experimental conditions) show a strong negative linear correlation ($r = -0.741$, $P = 0.0016$), indicating that decreased anisotropy (increased IRE1 oligomerization) is associated with increased cytoplasmic TXNIP localization. **(b)** Spearman correlation analysis of ranked TXNIP fluorescence versus ranked Apollo-IRE1 anisotropy values. Single-cell measurements ($N$ > 10,000 cells) reveal a moderate, negative, monotonic correlation ($\rho = -0.435$, $P < 0.0001$), confirming the inverse relationship between IRE1 activation state and TXNIP subcellular distribution at the single-cell level. **(c)** Summary table of correlation analyses between TXNIP fluorescence and Apollo-IRE1 anisotropy showing both per-replicate means and raw single-cell measurements for Pearson and Spearman correlations.

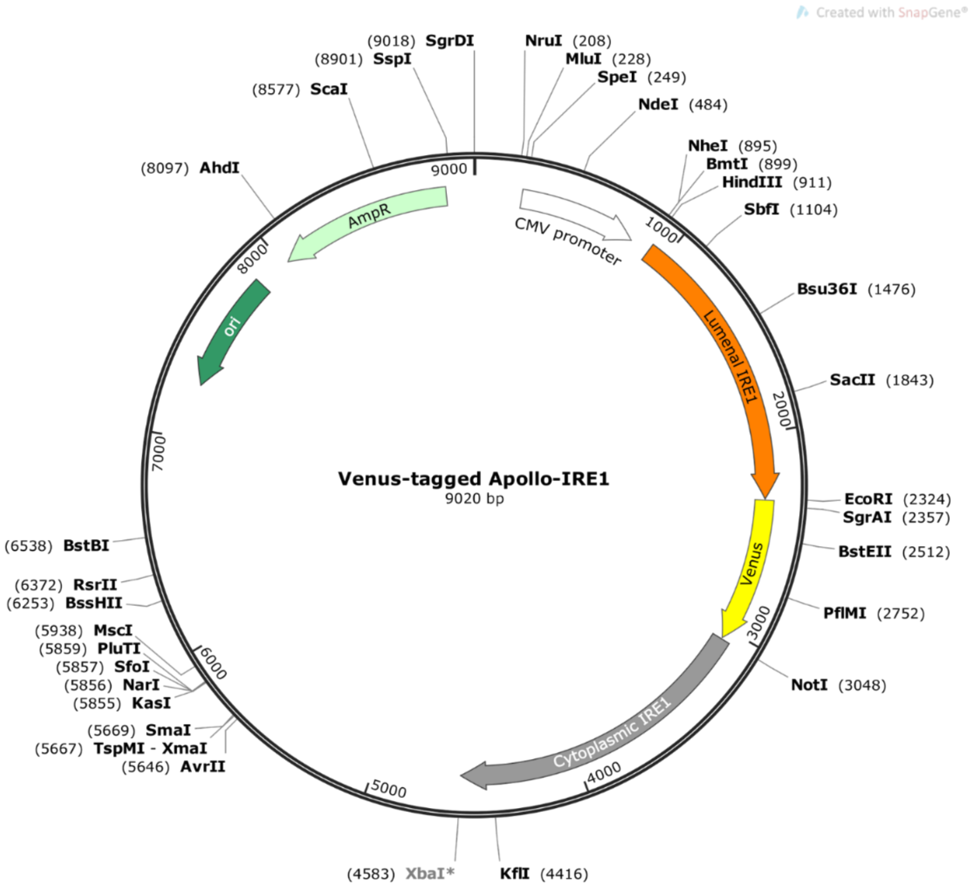


**Figure S3. Plasmid map of mVenus-tagged Apollo-IRE1 construct.** Schematic representation of the Apollo-IRE1 expression vector (9020 bp). The construct was generated by cloning the IF2L mutant of human IRE1α (WLLI359-362 to GSGS) into a pcDNA3.1(+) backbone, adapted from Karagöz et al.[24] The mVenus fluorescent protein is inserted intrasequence between the lumenal domain and the cytoplasmic kinase/RNase domain, positioned at the cytoplasmic face of the transmembrane domain. Key features include: CMV promoter for mammalian expression, ampicillin resistance gene (AmpR) for bacterial selection, and a bacterial origin of replication (Ori).

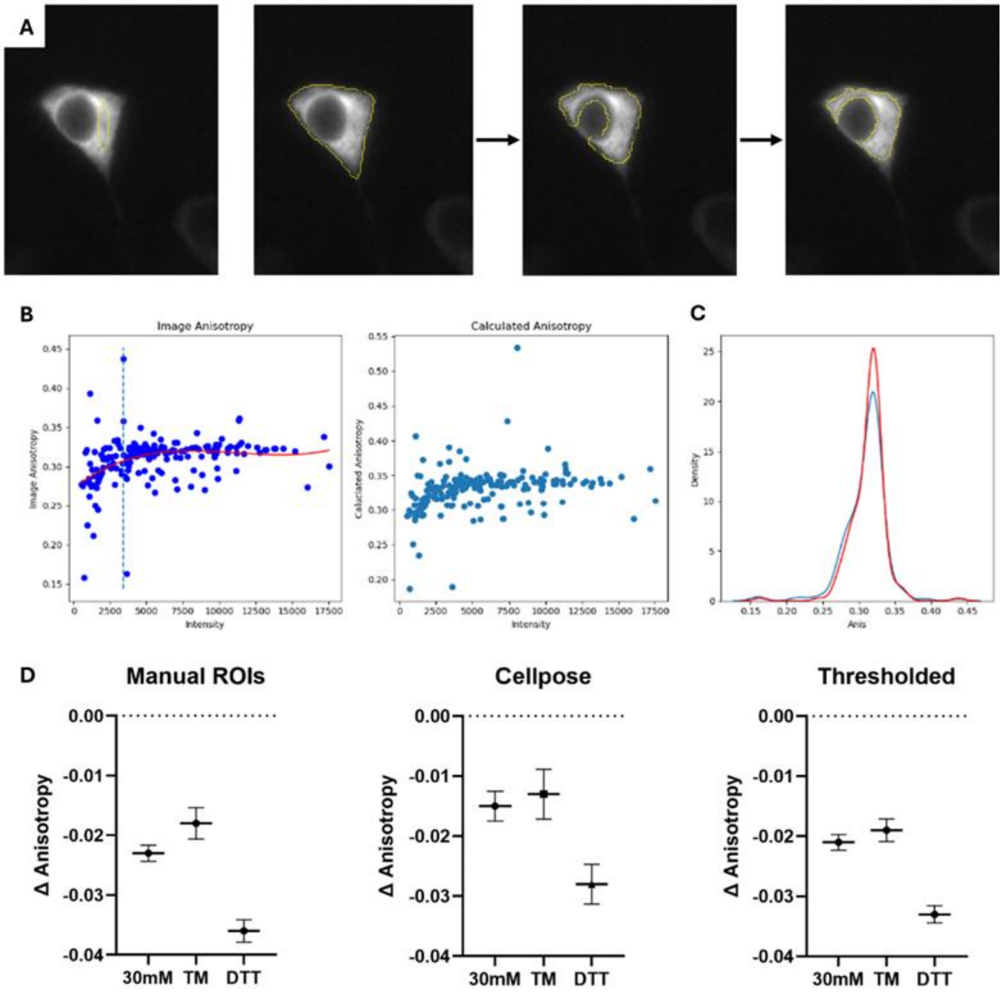


**Figure S4. Development and validation of automated high-throughput image analysis pipeline. (a)** Progression of region of interest (ROI) construction methods: from left to right: manual ROI drawn by hand; automated ROI generated by the Cellpose deep learning algorithm; ROI refined by Otsu automatic thresholding; final ROI after shrinking by 2 pixels in all directions to exclude nuclear regions. **(b)** Scatter plots of single-pixel anisotropy versus fluorescence intensity values demonstrating intensity-dependent variance in anisotropy measurements. The left panel shows raw image anisotropy, and the right panel shows calculated anisotropy after applying an intensity threshold. Pixels below 1500 intensity units (vertical dashed line) show higher anisotropy variance and are excluded from analysis. **(c)** Histogram distributions of anisotropy values before (left) and after (right) removal of low-intensity pixels demonstrate improved measurement precision following intensity thresholding. **(d)** Comparison of Apollo-IRE1 anisotropy measurements (Δ Anisotropy relative to control) across three ROI construction methods: manual ROIs, Cellpose-generated whole-cell ROIs, and thresholded/shrunken ROIs with nuclear exclusion. All three methods yield consistent results for ER stress treatments (30 mM glucose, tunicamycin (TM), and DTT), validating the automated pipeline. Data are presented as mean ± S.E.M.
